# Quantitative proteome and PTMome analysis of *Arabidopsis thaliana* root responses to persistent osmotic and salinity stress

**DOI:** 10.1101/2020.12.28.424236

**Authors:** MC. Rodriguez, D Mehta, M Tan, RG Uhrig

## Abstract

Abiotic stresses such as drought result in large annual economic losses around the world. As sessile organisms, plants cannot escape the environmental stresses they encounter, but instead must adapt to survive. Studies investigating plant responses to osmotic and/or salt stress have largely focused on short-term systemic responses, leaving our understanding of intermediate to longer-term adaptation (24 h - days) lacking. In addition to protein abundance and phosphorylation changes, evidence suggests reversible lysine acetylation may also be important for abiotic stress responses. Therefore, to characterize the protein-level effects of osmotic and salt stress, we undertook a label-free proteomic analysis of *Arabidopsis thaliana* roots exposed to 300 mM Mannitol and 150 mM NaCl for 24 h. We assessed protein phosphorylation, lysine acetylation and changes in protein abundance, detecting significant changes in 245, 35 and 107 total proteins, respectively. Comparison with available transcriptome data indicates that transcriptome- and proteome-level changes occur in parallel, while PTMs do not. Further, we find significant changes in PTMs and protein abundance involve different proteins from the same networks, indicating a multifaceted regulatory approach to prolonged osmotic and salt stress. In particular, we find extensive protein-level changes involving sulphur metabolism under both osmotic and salt conditions as well as changes in protein kinases and transcription factors that may represent new targets for drought stress signaling. Collectively, we find that protein-level changes continue to occur in plant roots 24 h from the onset of osmotic and salt stress and that these changes differ across multiple proteome levels.

## INTRODUCTION

Environmental conditions contributing to abiotic stresses such as drought and salinity result in large annual economic losses around the world. Between 2005 and 2015, drought was responsible for 30 % of agricultural loss, totaling to USD 19.5 billion (Food and Agriculture Organization of the United Nations (FAO), 2017). The combined result of human activities and climate change represent two of the major environmental factors currently threatening global food security (Ghosh and Xu, 2014), with rising global temperatures extending drought periods and lengthening plant drought recovery time (Schwalm et al., 2017). More frequent and severe droughts have left farmers to depend on progressively unsustainable irrigation and fertilization practices to maintain crop yield, placing further stress on drainage patterns and salinity levels of farmland (Kaushal et al., 2018). Both drought and salinity stress create a hyperosmotic condition typically referred to as osmotic stress, while increased salinity has the additive effect of concurrently increasing ionic strength. These osmotic and ionic effects massively impede the ability of roots to take up water, forcing molecular, cellular and/or morphological changes in order to adapt and survive.

All plant organs must adapt to drought and salt stress; however, roots are at the forefront and are thus responsible for adjusting their molecular program as necessary to ensure whole plant survival. In particular, under water-limiting conditions, roots restructure their investment from lateral root formation to axile roots to forage deeper soil layers (Paez-Garcia et al., 2015). For example, overexpression of the cell wall expansion protein EXPA5 (AT3G29030), stimulates root cell elongation by lengthening the root tip rather than increasing cell number (Xu et al., 2014). While DEEPER ROOTING 1 (DRO1; AT1G72490), a rice quantitative trait locus, is capable of increasing root angle in response to gravity, creating a steeper and deeper root system architecture, in addition to a higher photosynthetic capability, less wilting, and a >30 % increase in filled grain under severe drought (Uga et al., 2013). Thus, root system changes help define the growth and structure of aboveground biomass.

Within all plant cells, proteins are at the heart of sensing and initiating signaling events that then launch stress-adaptive responses. While the majority of our understanding of system drought responses has been driven by transcriptomic responses (Kreps et al., 2002; Kilian et al 2007; Rest et al., 2016; Yin et al., 2019), integrating this knowledge with a protein-level understanding of a plant’s environmental stress response and adaptation over time through quantitative proteomics, offers opportunities for both fundamental understanding and biotechnological application.

Osmotic and salt stress have a major influence over crop productivity worldwide (Ashraf and Akram, 2009), making it essential that we continue to refine our understanding of how plants respond to these perturbations in order to breed and/or engineer climate resilient crops. Emphasizing this need are the multiple studies that have examined the effects of osmotic and/or salt stress across a range of photosynthetic eukaryotes using both global and targeted analyses to understand the range of molecular, biochemical, physiological and morphological responses these stresses elicit (Zhang et al., 2012). This compendium of osmotic and salt stress data encompass a vast multitude of plant systems, time-frames and applied technologies, ranging from minutes to days of stress exposure (Jiang et al., 2007), followed by transcriptomic (Kilian et al., 2007; Kreps et al., 2002; Rest et al., 2016; Yin et al., 2019), proteomic (Jiang et al., 2007; Koh et al., 2015; Luo et al., 2015; Zhang et al., 2012) and metabolomic (Shelden et al., 2016) analyses to assess how the root environment responds to either abiotic stress. However, given the large variability in sampling time frames and root growth approaches, coupled with: 1) limited in-parallel examination of multiple stresses and proteoforms within the same study, 2) established disconnects between temporal changes in transcripts and proteins (Baerenfaller et al., 2012; Petricka et al., 2012; Reiland et al., 2009; Seaton et al., 2018; Uhrig et al., 2019), and 3) limited investigations of the role post-translational modifications (PTMs) play in root osmotic and salt adaptation (Koh et al., 2015; Wu et al., 2016; Zhu, 2016), further quantitative proteomic analysis are required.

PTMs are a pillar of properly functioning eukaryotic cells. Of the known PTMs, protein phosphorylation and lysine acetylation are the two most abundant (Choudhary et al., 2009; Henriksen et al., 2012; Hartl et al., 2017; Sharma et al., 2014; Uhrig et al., 2019; Weinert et al., 2011), with plants maintaining a comparable phosphoprotein landscape to humans (PhosPhat, http://phosphat.uni-hohenheim.de/; (Heazlewood et al., 2008), ATHENA; http://athena.proteomics.wzw.tum.de/; (Mergner et al., 2020); Plant PTM Viewer, https://www.psb.ugent.be/webtools/ptm-viewer/; (Willems et al., 2019). More recently, global analysis of protein acetylation in Arabidopsis (Finkemeier et al., 2011; Hartl et al., 2017; Liu et al., 2018; Uhrig et al., 2019; Wu et al., 2011) and multiple crop species (Guo et al., 2020; Nallamilli et al., 2014; Singh et al., 2020; Smith-Hammond et al., 2014; Walley et al., 2018; Zhou et al., 2018; and Xue et al., 2018) have similarly identified thousands of lysine acetylation events occurring on proteins with diverse functions and subcellular localizations. These two highly abundant PTMs have also been found to intersect on proteins involved in the same cellular processes (Rao et al., 2014; Soufi et al., 2012; Tran et al., 2012; Uhrig et al., 2019) in addition to occurring on different proteins involved in the same metabolic pathways (Uhrig et al., 2019), suggesting protein phosphorylation and lysine acetylation are functionally intertwined on multiple scales.

To better understand osmotic and salt stress responses in roots, we endeavored to examine the intermediate-term responses (24 h) of *Arabidopsis thaliana* (Arabidopsis) roots subjected to either 300 mM Mannitol or 150 mM NaCl. Then using label-free quantitative (LFQ) proteomics, we quantified the changes in the Arabidopsis root proteome, phosphoproteome, and acetylome using an established sequential quantitative proteomic workflow to facilitate the simultaneous assessment of multiple proteoforms in the same study. Our findings demonstrate how protein abundance and PTMs change relative to one another and to the transcriptome, in addition to understanding intersections within, and between, each abiotic stress condition to reveal osmotic and/or salt-specific responses.

## RESULTS

### Quantification and localization of mannitol- and salt-induced protein phosphorylation, lysine acetylation and abundance changes

Twenty-one day old Arabidopsis seedlings grown in liquid MS media were subjected to either osmotic (300 mM mannitol)- or salt (150 mM NaCl)-stress for 24 h starting and ending at zeitgeber time 3 (ZT 3) and assessed for quantifiable changes in protein phosphorylation and/or lysine acetylation status as well as protein abundance (Figure 1A; Supplemental Table 1-3). We quantified a total of 3,681 phosphorylated peptides and 266 acetylated peptides across 1,291 and 201 proteins, respectively, with a PTM localization ≥ 0.75 (Supplemental Table 2-3), while a total of 3,128 proteins were quantified for changes in abundance (Supplemental Table 1). Of these, 3681 phospho- and 266 acetyl-peptides corresponding to 1291 and 201 proteins, respectively, exhibited significant changes under osmotic stress (*pval* ≤ 0.05; FC ≥ 1.5), while 3681 phosphorylated and 266 acetylated peptides found on 1291 and 201 proteins, respectively, exhibited significant change (*pval* ≤ 0.05; FC ≥ 1.5) under salt stress (Figure 1B-D; Table 1; Supplemental Table 2-3). A total of 32 and 86 proteins exhibited significant changes in abundance (*pval* ≤ 0.05; FC ≥ 1.5) under osmotic and salt stress (Figure 1B-D; Table 1; Supplemental Table 1).

**Figure 1:**
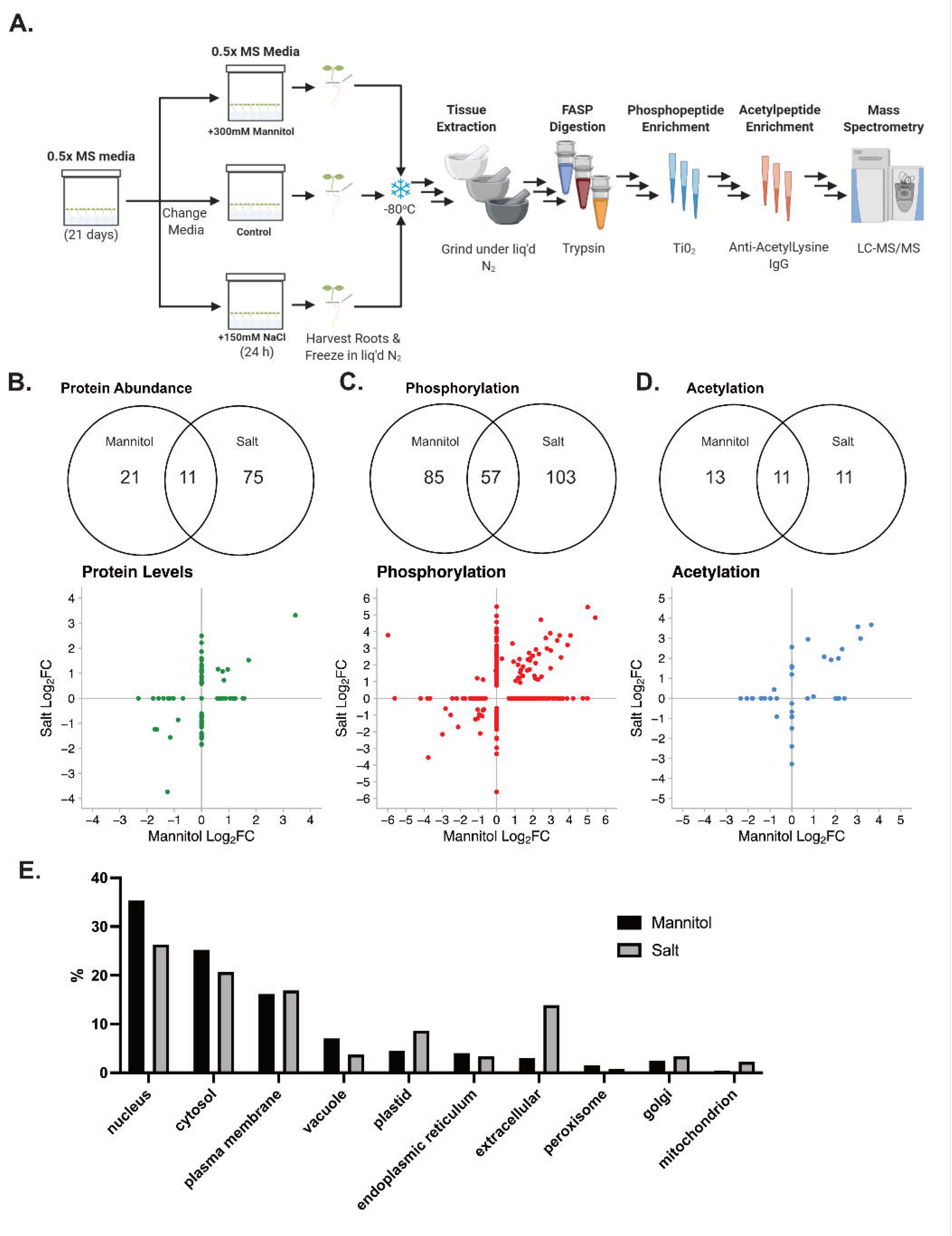
Proteome and PTMome changes under osmotic or salt stress. (A) Experimental workflow (B-D) Venn diagram and Log_2_ fold-change (FC) scatterplots for significantly changing (B) acetylome, (C) phosphoproteome and (D) proteome (unmodified proteins) across biological replicates (n=3). PTM-levels are plotted based on the median Log_2_FC of all modified peptides for each protein. (E) Subcellular localization distribution of significantly changing proteins in abundance or PTM-level (*pval* ≤ 0.05; FC ≥ 1.5).

**Table 1:** Summary results of proteomic and PTMomics analyses of Arabidopsis roots after 24 h of salt and osmotic stress (n=3). The last two columns depict the number (%) of significantly changing PTMs occurring on proteins either ^ii^exhibiting or ^i^not exhibiting a corresponding change in protein abundance is shown.

| Stress | Proteome Assessed | *Quantified |  | **Significantly Changing under Stress |  | <sup>i</sup>Number (%) of Sig PTM Quantified not Exhibiting Proteome Changes | <sup>ii</sup>Number (%) of Sig PTM Quantified Exhibiting Proteome Changes |
| --- | --- | --- | --- | --- | --- | --- | --- |
|  |  | Proteins | Peptides | Proteins | Peptides | Proteins | Proteins |
| 300mM Mannitol (Osmotic) | Acetylproteome (Ac) | 201 | 266 | 24 | 25 | 17 (71%) | 0 (0%) |
|  | Phosphoproteome (P) | 1291 | 3681 | 142 | 175 | 31 (21.9%) | 0 (0%) |
|  | Proteome (TP) | 3128 | n/a | 32 | n/a | n/a | n/a |
| 150mM NaCl (Salinity) | Acetylproteome (Ac) | 201 | 266 | 22 | 25 | 13 (59%) | 0 (0%) |
|  | Phosphoproteome (P) | 1291 | 3681 | 160 | 203 | 39 (24.4%) | 0 (0%) |
|  | Proteome (TP) | 3128 | n/a | 86 | n/a | n/a | n/a |

\*Quantified PTM (Ac & P) = Localization $\geq 0.75$
\*Quantified Proteome (TP) = min 2 peptides per protein
\*\*Significantly changing under stress (Ac, P & TP) = $p\text{-val} \leq 0.05$ and $\geq 1.5$ fold-change in PTM abundance

Next, we compared the Log_2_ fold-change (log_2_FC) distribution of significantly changing proteins and PTMs between osmotic- and salt-treated roots (Figure 1B-D). Analysis of proteins exhibiting a significant change in either abundance, phosphorylation or lysine acetylation found a total of 11 (11.5 %), 57 (30.3 %) and 11 (45.8 %) proteins overlapping between stress conditions (Figure 1B-D). Lastly, to understand where within the plant cell environment intermediate-term changes in the proteome are occurring, we obtained the predicted subcellular localization for all proteins exhibiting a significant change in PTM status or abundance under either osmotic- or salt-stress using SUBAcon (SUBA4; https://suba.live/; Hooper et al., 2014). Here, we found differences in the subcellular distribution of significantly changing proteins between each treatment (Figure 1E), with both mannitol and salt predominantly inducing changes in cytosolic, nuclear, and plasma membrane proteins. Salt also elicited changes in the greater numbers of plastidial and extracellular proteins, while mannitol had more changes in proteins localized to the vacuole (Figure 1E).

### Overlap between changing PTMome and proteome is limited

Often, PTMs are quantified independent of assessing changes in proteome abundance. Therefore, reported changes in PTM levels may also be due to fluctuations in protein abundance. To better understand this here, we examined proteins exhibiting PTM changes for concurrent changes in protein abundance. Under osmotic stress 31 (21.9 %) and 17 (71 %) proteins exhibiting significant changes in either phosphorylation or lysine acetylation were also quantified at the proteome level to not be changing in abundance. No PTM modified proteins concurrently exhibited a significant change in PTM and protein abundance (Table 1). Similarly, under salt stress, 39 (24.4 %) and 13 (59 %) phosphorylated and acetylated proteins, respectively, were quantified in our proteome data to not be changing (Table 1). Our concurrent assessment of proteome and PTM abundance changes provides increased confidence that observed phosphorylation and lysine acetylation changes are related to PTM-level changes, and not changes in proteome abundance. Moreover, since assessing changes in PTMs requires enrichment of modified peptides, it is likely that the remainder of PTM modified proteins not observed in our protein abundance measurements are simply below detection.

### Gene Ontology (GO) analysis of proteome and PTM fluctuations reveals differences between osmotic- and salt-induced stress

To better understand what cell processes are changing at the proteome-level after 24 h of osmotic or salt stress, we performed a gene ontology (GO) enrichment analysis for biological process GO terms (Supplemental Table 4). In response to salt stress, we found a number of significantly enriched GO terms amongst proteins exhibiting a change in abundance. These include: response to osmotic stress (GO:0006970, sulfate reduction (GO:0019419), S-glycoside metabolic processes (GO:0016143), glucosinolate metabolic processes (GO:0019757), in addition to organonitrogen metabolism (GO:1901564), translation (GO:0006412), and translational elongation (GO:0006414) (Table 2). Similarly, under osmotic stress we see enrichment of organonitrogen compound catabolism (GO:1901565), sulfur compound metabolic processes (GO:0006790) and S-glycoside metabolic processes (GO:0016143) amongst the significantly changing proteins (Supplemental Table 4). At the phosphoprotein level, salt stress induced changes in a number of biological processes. Here we saw significant enrichment of cyclic nucleotide metabolic processes (GO:0009187), protein phosphorylation (GO:0006468) and cell surface receptor signaling pathway (GO:0007166). Conversely, osmotic stress conditions saw the enrichment of cell division (GO:0051301), cell cycle (GO0007049) and cell cycle processes (GO:0022402). Next, we performed a protein domain analysis using Thalemine (https://bar.utoronto.ca/thalemine/). Here, we found an enrichment of a number of protein domains amongst the proteins significantly changing in abundance under salt stress (Supplemental Table 4). In particular, multiple chaperone and chaperonin domains as well as thioredoxin domains in addition to glycoside hydrolase superfamily, which supports our GO findings of augmented S-glycoside metabolic processes (GO:0016143) in salt stressed roots.

**Table 2:** Enriched GO terms for significant protein abundance changes upon 24 h of 150 mM salt (NaCl) treatment

| GO Term | GO ID | pvalue | BH pvalue |
| --- | --- | --- | --- |
| organonitrogen compound metabolic process | GO:1901564 | 3.63E-06 | 6.51E-04 |
| S-glycoside metabolic process | GO:0016143 | 7.48E-06 | 0.001207625546 |
| translational elongation | GO:0006414 | 2.01E-05 | 0.002457265017 |
| glycosyl compound biosynthetic process | GO:1901659 | 2.20E-05 | 0.00257978525 |
| sulfate reduction | GO:0019419 | 1.42E-04 | 0.01381235311 |
| glycosinolate metabolic process | GO:0019757 | 1.44E-04 | 0.01381235311 |
| glycosinolate biosynthetic process | GO:0019758 | 1.96E-04 | 0.01814212421 |
| translation | GO:0006412 | 2.10E-04 | 0.01882462124 |
| regulation of endopeptidase activity | GO:0052548 | 4.64E-04 | 0.03122389969 |
| response to osmotic stress | GO:0006970 | 4.81E-04 | 0.03157020814 |
| organonitrogen compound biosynthetic process | GO:1901566 | 6.49E-04 | 0.03968781631 |
| protein dephosphorylation | GO:0006470 | 7.03E-04 | 0.04111057319 |

| glycosyl compound metabolic process | GO:1901657 | 8.62E-04 | 0.04916442018 |
| --- | --- | --- | --- |
| cellular amino acid metabolic process | GO:0006520 | 8.77E-04 | 0.04916442018 |
| Protein Domain Name | AGI | Domain | BH pvalue |
| Chaperonin Cpn60/TCP-1 family | AT1G24510,AT3G02530,AT3G03960,AT3G11830,AT5G20890,AT5G26360,AT5G56500 | IPR002423 | 0.0001977420788 |
| Heat shock protein 70 family | AT3G09440,AT4G37910,AT5G02500,AT5G09590,AT5G28540,AT5G42020 | IPR013126 | 0.001032156851 |
| Pyridoxal phosphate-dependent transferase | AT1G23310,AT1G43710,AT1G80600,AT2G13360,AT2G20610,AT3G01120,AT4G32520,AT4G35630,AT5G17330,AT5G26780,AT5G46180 | IPR015424 | 0.001649480794 |
| Thioredoxin-like superfamily | AT1G02920,AT1G09640,AT1G35620,AT1G45145,AT1G49860,AT1G57720,AT1G62180,AT1G65980,AT1G75270,AT2G47470,AT3G09270,AT4G02520,AT4G04610,AT4G04950,AT4G21990,AT4G27080,AT5G08290,AT5G47890 | IPR036249 | 0.002955649486 |
| Glutathione S-transferase, C-terminal | AT1G02920,AT1G09640,AT1G49860,AT1G57720,AT3G09270,AT4G02520 | IPR004046 | 0.003395888372 |
| NAD(P)-binding domain superfamily | AT1G04410,AT1G08200,AT1G13440,AT1G15950,AT1G31230,AT1G53240,AT1G63000,AT1G64190,AT1G67730,AT1G79530,AT3G07690,AT3G15290,AT3G19450,AT3G25530,AT3G58610,AT3G61220,AT4G34230,AT5G08300,AT5G14780,AT5G28840,AT5G43330 | IPR036291 | 0.003434118093 |
| Hyaluronan/mRNA-binding protein | AT4G16830,AT4G17520,AT5G47210 | IPR006861 | 0.004099704067 |
| Aminoacyl-tRNA synthetase, class I, conserved site | AT1G14610,AT1G25350,AT1G66530,AT4G10320,AT4G13780 | IPR001412 | 0.004388306174 |
| Six-bladed beta-propeller, TolB-like | AT1G21670,AT1G74020,AT2G01410,AT2G47390,AT3G57020,AT5G39970 | IPR011042 | 0.01776705313 |
| Glycoside hydrolase superfamily | AT1G47600,AT1G51470,AT3G13750,AT3G18080,AT3G52840,AT3G60130,AT4G19810,AT5G07830,AT5G08380,AT5G26000,AT5G40390,AT5G49360,AT5G56870,AT5G63800 | IPR017853 | 0.02231435074 |
| Histidine kinase/HSP90-like ATPase | AT2G04030,AT3G07770,AT4G24190,AT5G56010,AT5G56030 | IPR003594 | 0.03531610162 |
| 6-phosphogluconate dehydrogenase-like, C-terminal domain superfamily | AT1G64190,AT3G07690,AT3G15290,AT3G25530,AT3G58610 | IPR008927 | 0.03531610162 |

### Comparative analysis of proteome and transcriptome changes reveals osmotic and salt-response disconnects

It is well established that transcriptome and proteome changes are often disconnected in their temporal peak expression patterns (Baerenfaller et al., 2012; Graf et al., 2017; Seaton et al., 2018). Therefore, using publicly available microarray transcriptome data from Arabidopsis plants subjected to analogous stress treatments under similar growth conditions (Kilian et al., 2007), we performed a comparative analysis to better understand the temporal dynamics of the transcripts that correspond to proteins exhibiting phosphorylation, lysine acetylation or abundance changes upon 24 h of osmotic- or salt-stress. Despite being comparable, some differences exist between our experimental conditions and that of Kilian and colleagues (2007). These include: the use of a 12:12 photoperiod, 21 day old plants and stress application starting at zeitgeber (ZT) 3 in our study versus a 16:8 photoperiod, 18 day old plants and an unknown stress application start time by Kilian and colleagues (2007). Using this transcriptome data, we sought to cluster corresponding transcripts based on their expression dynamics and investigate how these dynamics were reflected at the level of protein abundance as well as lysine acetylation or phosphorylation status (Figure 2–3; Supplemental Table 5).

**Figure 2:**
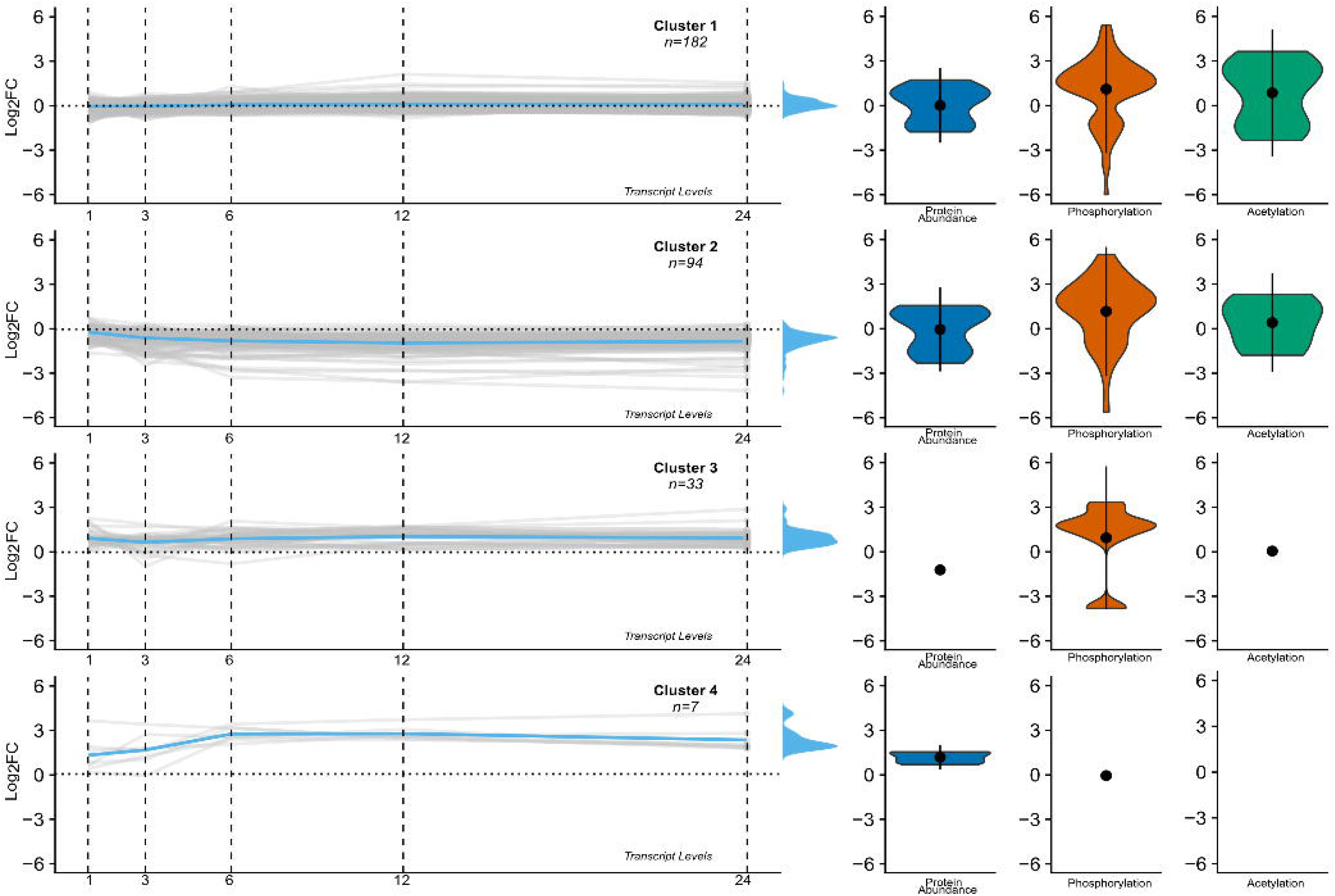
Comparison of transcript-level changes with corresponding protein abundance, phosphorylation, and lysine acetylation changes upon 24 h of osmotic stress. Transcript-level changes corresponding to significantly changing proteins (either abundance or for each PTM) in our study were calculated from previously described microarray experiments (KIllan et al., 2007) and clustered by longitudinal shape.

**Figure 3:**
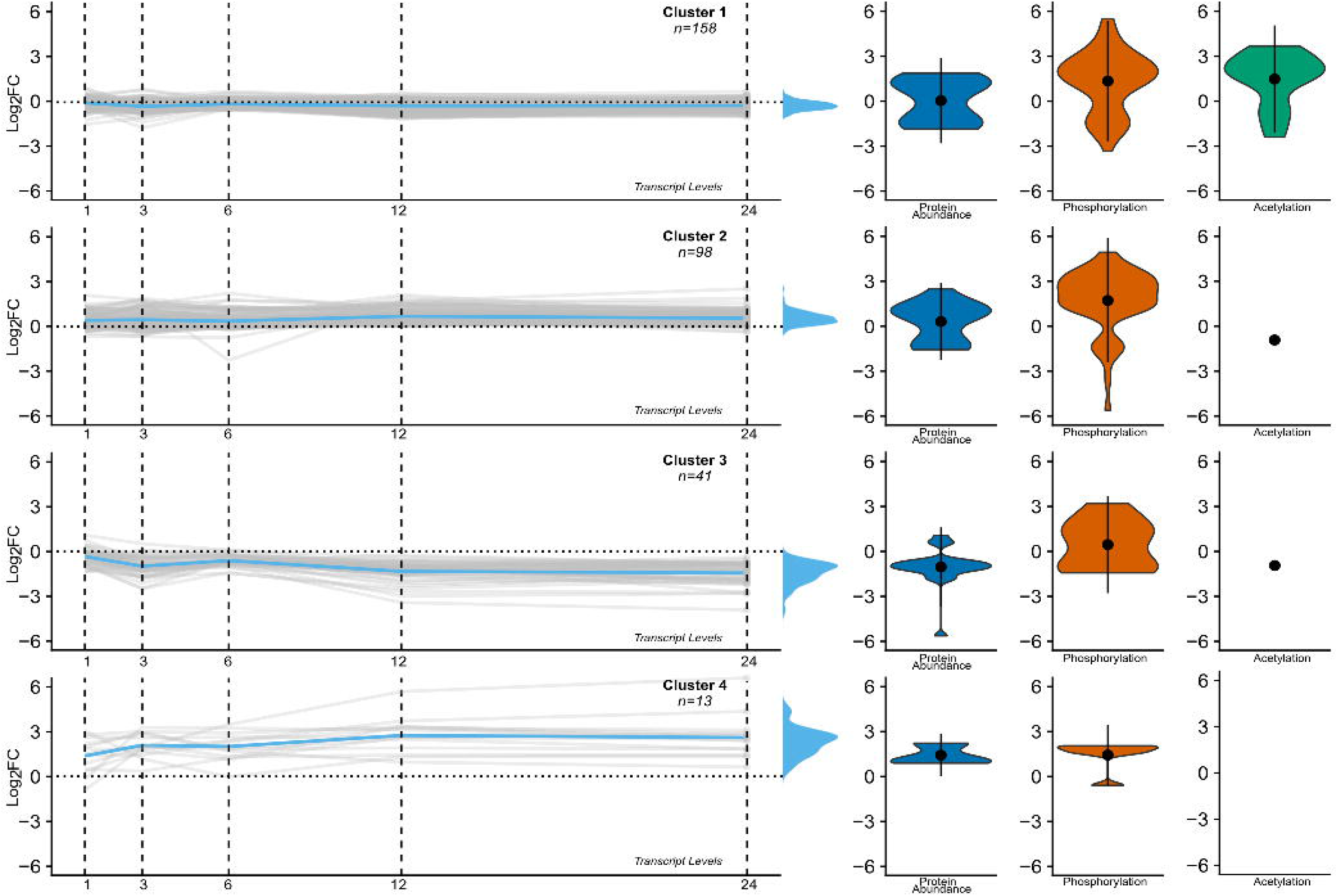
Comparison of transcript-level changes with corresponding protein abundance, phosphorylation, and lysine acetylation changes upon 24 h of salt stress. Transcript-level changes corresponding to significantly changing proteins in our study (either based on abundance or for each PTM) were calculated from previously described microarray experiments (Killan et al., 2007) and clustered by longitudinal shape.

Under both osmotic- and salt-stress conditions, the majority of proteins exhibiting a significant change in abundance, phosphorylation and/or lysine acetylation 24 h post-stress maintained a minimal log_2_FC transcriptional change (Cluster 1 - Figure 2–3). We also saw protein clusters emerge under both osmotic- and salt-stress whereby an increasing trend in their transcript abundance from stress onset (0 h) to 24 h post-stress paralleled our observed end-point proteome abundance changes at 24 h (Cluster 2 - Figure 2–3). However, this was not case for protein phosphorylation and acetylation events, which did not parallel transcriptome changes in either the osmotic or salt stress treatments (Figure 2–3). By clustering the significantly changing proteome and PTMome based on their corresponding transcriptional expression profile from a comparably acquired, publicly available dataset, we find that transcript changes are generally a good indicator of proteome changes 24 h post-osmotic or -salt stress initiation in Arabidopsis roots. Conversely, we find that transcriptional changes are a poor indicator of PTM-level changes. Overall, our data finds that osmotic and salt-stress induced proteomes, when clustered by transcriptional change profile (0 h to 24 h), reveals key differences in how Arabidopsis roots respond to these stresses after 24 h.

### Network analysis of proteins exhibiting osmotic- and salt-induced changes in PTM status and/or abundance reveals how the proteomes intersect

Next, using a combination of STRING-DB association network analyses (https://string-db.org/) and SUBA4 (https://suba.live/) protein subcellular localization information, we assembled subcellular localization-resolved protein association networks to elucidate the degree of interconnectedness between osmotic- and salt-induced PTM and protein abundance changes (Figure 4–5). Proteins with an edge score threshold of 0.5 or below were excluded from visualization. Here, we find that changes in the PTM status and/or protein abundance of metabolic proteins were most common between both osmotic and salt stress conditions (Figure 4–5). This is highlighted by proteins involved in both the transport and assimilation of sulfur as well as primary metabolism (Figure 4–5). Similarly, changes in transcription and protein translation were also common between osmotic and salt treatments. Lastly, we see a number of known drought responsive proteins to be changing in both our osmotic and salt condition, indicating that our treatment regimes elicited the expected responses.

**Figure 4:**
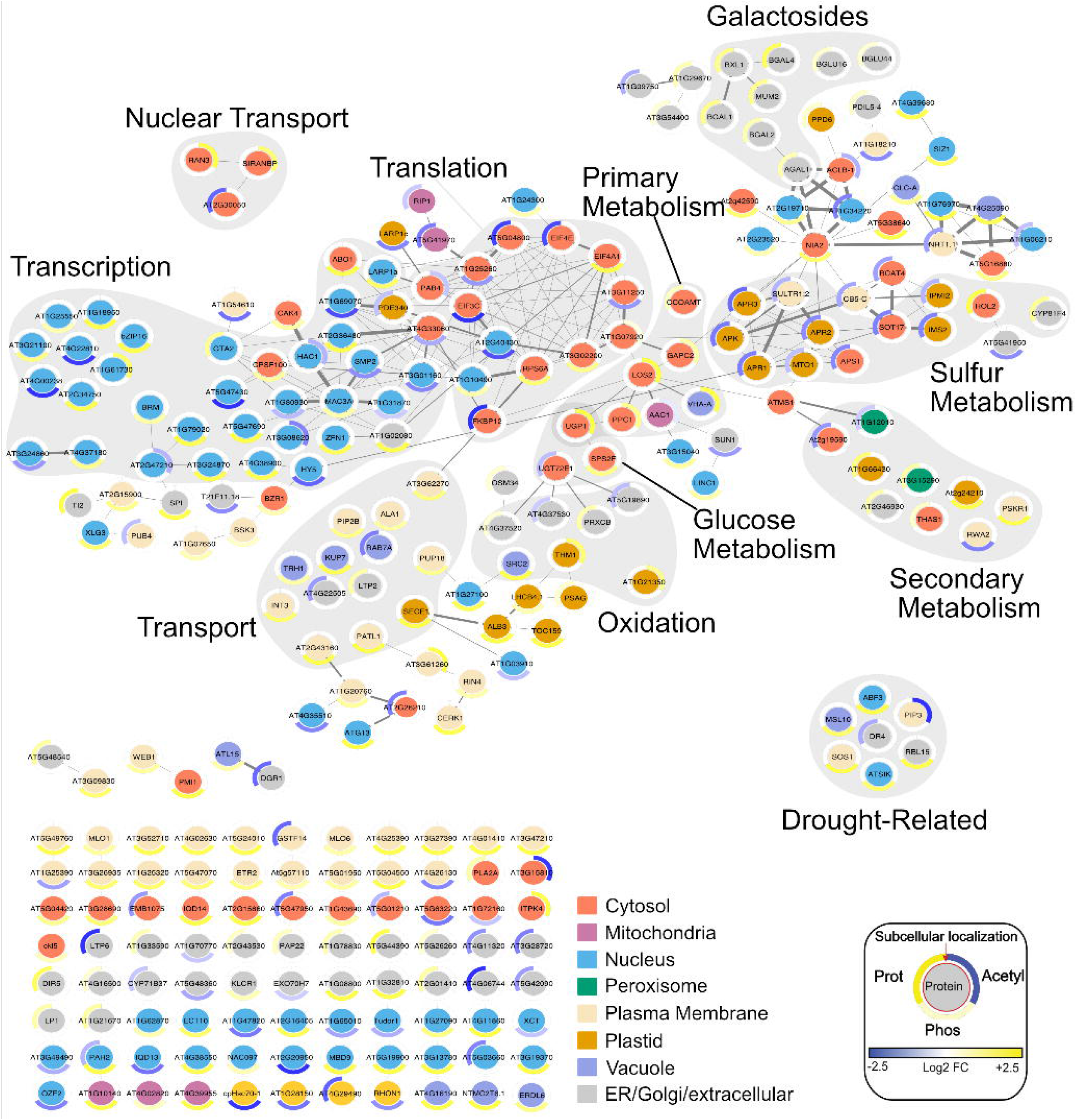
Association network of Arabidopsis root under 24 h 150 mM salt (NaCl). STRING database association network shown for proteins significantly changing in their total protein, phosphorylation, or lysine acetylation status. The maximal log_2_FC change ranges from −2.5 (Max log_2_FC EN) to +2.5 (Max log_2_FC ED). Edge thickness equates to confidence in association. Inner circle represents subcellular localization (SUBA 4; https://suba.live/): cytosol (red), mitochondria (purple), nucleus (blue), peroxisome (green), plasma membrane (beige), plastid (amber), vacuole (lavender), endoplasmic reticulum/extracellular/Golgi (grey). Outer ring represents protein status: total protein change (1^st^ division), phosphorylation (2^nd^ division), lysine acetylation (3^rd^ division). Grey clusters depict proteins belonging to a common cellular process. Proteins with an edge score threshold of 0.5 or below were excluded from visualization.

**Figure 5:**
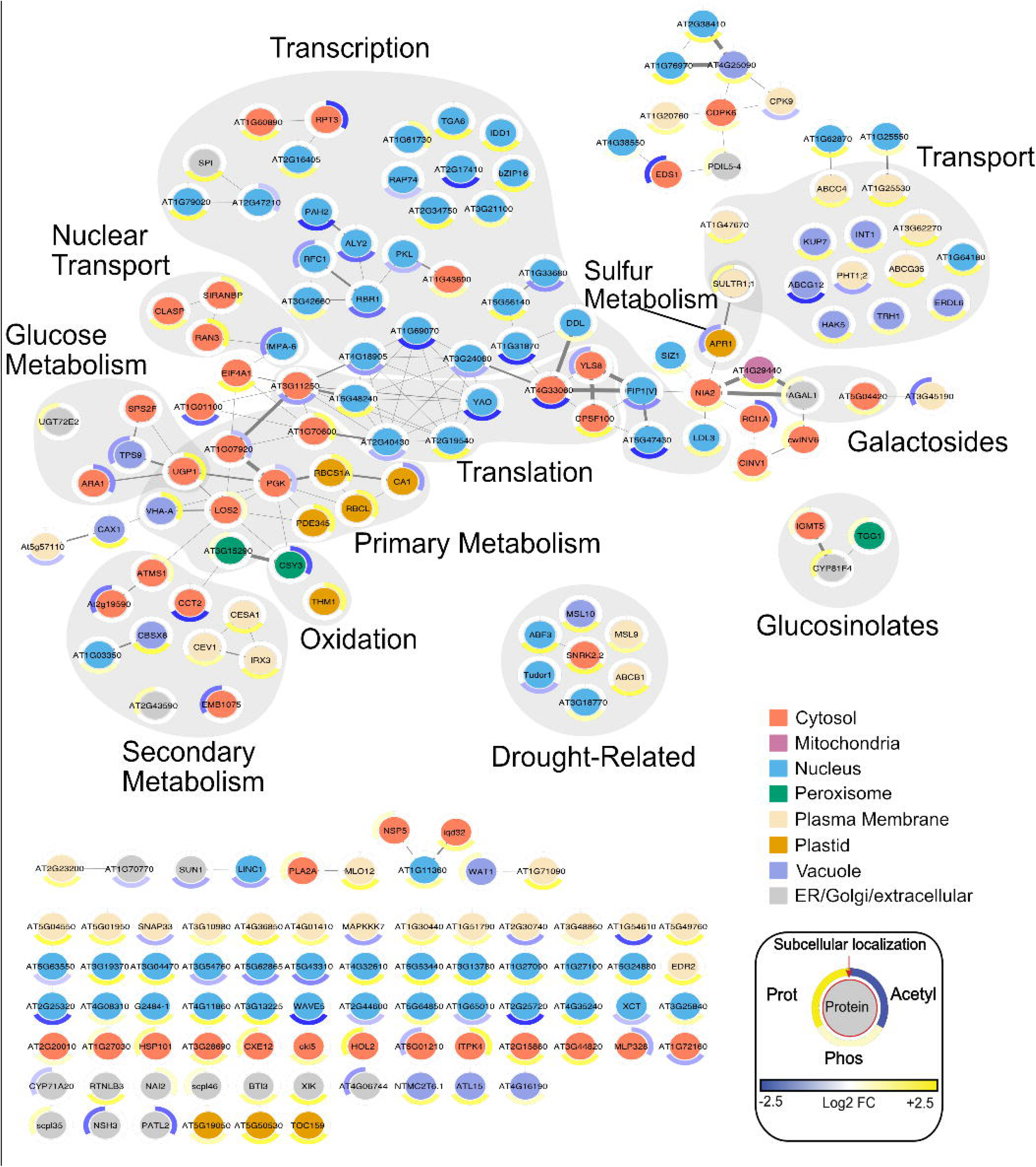
Association network of Arabidopsis root under 24 h 300 mM mannitol. STRING database association network shown for proteins significantly changing in their total protein, phosphorylation, or lysine acetylation status. The maximal log_2_FC change ranges from −2.5 (Max log_2_FC EN) to +2.5 (Max log_2_FC ED). Edge thickness equates to confidence in association. Inner circle represents subcellular localization (SUBA 4; https://suba.live/): cytosol (red), mitochondria (purple), nucleus (blue), peroxisome (green), plasma membrane (beige), plastid (amber), vacuole (lavender), endoplasmic reticulum/extracellular/Golgi (grey). Outer ring represents protein status: total protein change (1^st^ division), phosphorylation (2^nd^ division), lysine acetylation (3^rd^ division). Grey clusters depict proteins belonging to a common cellular process. Proteins with an edge score threshold of 0.5 or below were excluded from visualization.

Overall, our association networks further validate earlier GO enrichment analyses in addition to visualizing the strength of these relationships using compiled meta-association data between proteins. These networks also provided a means by which to integrate additional data types including: the type and degree of change in lysine acetylation, phosphorylation and/or abundance (outer ring), in addition to the subcellular localization of each protein within the plant cell (inner circle). Together, this data highlights the multi-dimensional relationships that exist between proteins exhibiting significant changes in either abundance or PTM status in response to osmotic and/or salt stress.

## DISCUSSION

### Benchmarking our data against known drought responsive proteins

The identification of the following proteins in our protein association networks suggests that our application of mannitol (300 mM) and salt (150 mM) was sufficient to elicit a stress response in 21 day old Arabidopsis plants. In our data, we find DROUGHT REPRESSED 4 (AtDR4; AT1G73330) a protease inhibitor protein that decreases in transcript abundance upon increasing drought (Frei dit Frey et al., 2010), to also decrease in protein abundance upon salt stress (Log_2_FC = −1.02). PATATIN-RELATED PHOSPHOLIPASE A (pPLA; AT2G26560) which is involved in modulating Arabidopsis water status (Yang et al., 2012) is up-regulated under both osmotic (Log_2_FC = 0.96) and salt (Log_2_FC = 1.15) conditions (Supplemental Table 1). From a PTM perspective, SALT-OVERLY-SENSITIVE 1 (SOS1; AT2G01980), a plasma membrane Na(+)/H(+) antiporter (Shi et al., 2000) and characterized salt tolerance protein (Ullah et al., 2016) that is activated by phosphorylation (Quintero et al 2011) increases in its phosphorylation status (Log_2_FC = 2.7) after 24 h of salt treatment. Further, osmotic stress results in elevated SNF1-RELATED PROTEIN KINASE 2.2 (SNRK2.2; AT3G50500) phosphorylation (Log_2_FC = 2.71). SNRK2.2 is a key activator of ABA signaling pathway that regulates responses such as stomatal closure (Kulik et al., 2011). We also see a decrease in the phosphorylation of TUDOR-SN PROTEIN 1 (TSN1; AT5G07350) under salt stress (Log_2_FC = − 0.96), which would suggest this is an activating PTM event, as TSN1 is known to regulate GIBBERELLIC ACID 20 OXIDASE 3 (GA20ox3; AT5G07200) expression under salt stress (Yan et al., 2014) (Supplemental Table 2). We further expand the role of TSN1 in this capacity beyond salt stress to also include osmotic stress, as we similarly see a decrease in phosphorylation (Log_2_FC = −0.95). Lastly, PLASMA MEMBRANE INTRINSIC PROTEIN 2;7 (PIP2;7; AT4G35100), an aquaporin known to be involved in osmoregulation during drought and salinity stress and to be extensively modified by multiple PTMs (Di Pietro et al., 2013), exhibited a decrease in its lysine acetylation status under salt stress (Log_2_FC = −3.28) (Supplemental Table 3). The fluctuation of these aforementioned proteins across the various proteomes we assessed provides confidence that our dataset contains novel protein-level insights into plant osmotic and salt stress responses. Further, the concerted changes in both the transcriptome and significantly changing proteome observed in both our osmotic or salt treatments, suggests our treatments are in line with previous osmotic and salt treatments, and demonstrates the utility of performing meta-analyses to more broadly understand and contextualize results.

### Protein lysine acetylation is up-regulated on core facets of glycolysis under both osmotic and salt stress conditions

Protein lysine acetylation occurring on primary glycolytic enzymes is almost universally up-regulated under both osmotic and salt stress (Figure 1B). For example, under both osmotic and salt treatments, key metabolic enzymes between glyceraldehyde 3-phosphate and oxaloacetate, such as GLYCERALDEHYDE-3-PHOSPHATE DEHYDROGENASE C2 (GAPC2; AT1G13440; NaCl_Log_2_FC = 1.19), PHOSPHOGLYCERATE KINASE 3 (PGK3; AT1G79550; MAN_Log_2_FC = −0.69), ENOLASE 2 (LOS2; AT2G36530; MAN_Log_2_FC = 0.74, NaCl_Log_2_FC = 2.95), and PHOSPHOENOLPYRUVATE CARBOXYLASE (PPC1; AT1G53310; NaCl_Log_2_FC = 1.60) all have upregulated lysine acetylation events. Similarly, UDP-GLUCOSE PYROPHOSPHORYLASE 1 (UGP1; AT3G03250; MAN_Log_2_FC = 3.02, NaCl_Log_2_FC =3.56) also shows an unidirectional up-regulation in acetylation status under both applied stresses (Figure 4–5; Supplemental Table 3).

Protein lysine acetylation has recently become established as a regulatory PTM in plants and other organisms (Finkemeier et al., 2011, Hartl et al., 2017; Zhao et al., 2010). Catalyzed by acetyltransferases, protein acetylation requires the primary metabolite acetyl-CoA. Acetyl CoA is a central intermediate metabolite between catabolic and anabolic pathways and as such, it is uniquely placed to exert either feedback or feedforward regulation through reversible protein acetylation. Interestingly, we see two key metabolic enzymes directly connected to acetyl-CoA metabolism to exhibit decreases in their acetylation status under osmotic or salt stress (Figure 4 and 5). These include: CITRATE SYNTHASE 3 (CSY3; AT2G42790; MAN_Log_2_FC = −2.06) and ATP-CITRATE LYASE B-1 (ACLB-1; AT3g06650; NaCl_Log_2_FC = −0.94). Multiple studies have found that protein N^ε^ acetylation is an abundant PTM across plants (De Boor et al., 2015; Finkemeier et al., 2011; Hartl et al., 2017; Li et al., 2018; Uhrig et al., 2019; Wu et al., 2011). In particular, glycolytic and specialized metabolic proteins have been found to be extensively acetylated under nitrogen stress in tea leaves (Jiang et al., 2018) as well as in a diel manner across a number of facets of plant metabolism in *Arabidopsis thaliana*, including N^ε^ acetylation of GAPC in roots (Uhrig et al., 2019). Here, our data suggest that N^ε^ acetylation also plays a role in salt and osmotic stress response in plants through primary metabolism. Interestingly, N^α^ protein acetylation has also been connected to drought resistance and is suggested to be ABA stress responsive (Linster et al., 2015; Linster et al., 2020). Since both N^α^ and N^ε^ protein acetylation rely on acetyl-CoA as an acetate donor, there likely exists higher-level intersections between these two forms of protein acetylation. Highlighting this possibility is the recent discovery of dual-specificity GNAT acetyltransferases in Arabidopsis (Bienvenut et al., 2020).

### Sulfur metabolism needed for short- and intermediate-term drought response

During drought stress, ABA is synthesized in the roots and then transported to the shoots to induce stomatal closure (Agurla et al., 2018). However, ABA biosynthesis in the roots requires sulfur. Plant root sulfate assimilation is primarily driven by ADENOSINE 5’PHOSPHOSULFATE REDUCTASE (APR), which reduces sulfate to sulfite followed by further reduction to sulfide by FERREDOXIN DEPENDENT SULFITE REDUCTASE (SiR) (Takahashi et al., 2011). Sulfide is then fixed by O-acetylserine(thiol)lyase into Cysteine (Cys) (Chan et al., 2013) to act as the sulfur donor for the sulfuration of molybdenum cofactor (Moco) sulfurase (Xiong et al., 2001), which is then necessary to initiate ABSCISIC ALDEHYDE OXIDASE 3 (AAO3) (Mendel, 2002; Schwarz and Mendel, 2006) and the ultimate production of ABA.

Studies have determined that Cys levels attune ABA biosynthesis (Batool et al., 2018; Cao et al., 2014), which in turn, limits sulfur assimilation at the level of APR (Takahashi et al., 2011). APR enzymatic activity is driven by the demand for sulfur, meaning that the uptake and assimilation of sulfur during drought is likely foundational to the short-term drought response. At 24 h of NaCl stress, we observed a number of key sulfur assimilating enzymes to decrease in their protein abundance, including: ATP SULFURYLASE 1 (APS1; AT3G22890; Log_2_FC = −1.43), APR1 (AT4G04610; Log_2_FC = −1.56), APR2 (AT1G62180; Log_2_FC = −1.39) and APR3 (AT4G21990; Log_2_FC = −1.80) and ADENOSINE 5’-PHOSPHOSULFATE KINASE (APK; AT2G14750 Log_2_FC = −1.60) (Figure 4–5; Supplemental Table 1). Furthermore, SULFATE TRANSPORTER 1;2 (SULTR1;2; AT1G78000), which mediates the uptake of the environmental sulfate by plant roots, is reduced in abundance (Log_2_FC = −0.90) under salt stress. Osmotic stress also induced sulfur related enzyme changes. SULTR1;1 exhibited an increase in protein abundance (Log_2_FC = 1.56), while APR1 was reduced in abundance similar to that observed under salt stress (Log_2_FC = − 1.15). This suggests that by 24 h post salt-stress initiation plants have already transitioned from utilizing short-term adaptations, to implementing longer term solutions.

Transcriptionally, APR is regulated by sulfur availability (Hirai et al., 2003), demand for Cys (Davidian and Kopriva, 2010), and also in response to environmental stressors such as high salt (Koprivova et al., 2008). Specifically, Koprivova and colleagues (2008) reported a 3-fold increase in APR mRNA after 5 h of 150 mM NaCl in Arabidopsis roots. Using the AtGenExpress Arabidopsis stress expression dataset (Kilian et al 2007), we calculated a peak expression of APR transcripts at 6 h post 150 mM NaCl stress initiation, which then was found to continually decrease through to 24 h (Kilian et al., 2007; Supplemental Table 5). Interestingly, an opposing trend in shoot APR expression is observed (Kilian et al., 2007). While APR in roots peaks soon after high salt exposure, APR in shoots remains low until eventually peaking at 24 h. This points to APR and sulfur driven drought-responses acting as a short-term responder in the roots and an intermediate responder in the shoots. Given this, a sharp increase in root APR levels may facilitate the sulfur assimilation required for ABA production, which then leads to prompt stomatal closure. A later peak in shoot APR may be useful for intermediate drought response mechanisms, such as reducing reactive oxygen species (ROS).

Salt stress is also known to increase the production of oxidative enzymes (e.g. NADPH oxidase), which are necessary for signaling and response during abiotic stress (Mittova et al., 2003), with subsequent reduction in ROS required as the salt stress persists (Hasegawa and Bressan, 2000). To mitigate the damaging effects of ROS, plants can produce glutathione (GSH), an antioxidant derived from Glu, Cys, and Gly. Stress-induced GSH production requires sulfur-containing Cys, with high levels of Cys found to enhance synthesis of GSH in plants (Takahashi et al., 2011). Demand for both, Cys and GSH, reciprocally regulates sulfur assimilation through APR (Davidian and Kopriva, 2010), with depletion of GSH pools under prolonged drought stress conditions driving the production of APR (Chan et al., 2013). Thus, upregulation of APR activity in the shoots may contribute to the production of antioxidants such as GSH. Further time-course experimentation is required to fully elucidate the interplay between salt-induced sulfur metabolism changes at the protein level in roots versus shoots, and is beyond the scope of this study.

### Both osmotic and salt stresses induce high-levels of UDP-glucose pyrophosphorylase (UGP1) lysine acetylation

We find that UGP1 (AT3G03250) maintained some of the largest increases in lysine acetylation, with salt (Log_2_FC = 3.58) and osmotic (Log_2_FC = 3.03) stress both eliciting large changes to UGP1 acetylation (Figure 2–3). UGP1 activity has previously been found to be regulated by PTMs. For example, phosphorylation by Per-Arnt-Sim (PAS) kinase downregulates UGP1 activity resulting in a net decrease of glycogen in yeast (Rutter et al., 2002). Analysis of available osmotic- and salt-stress induced transcript data found limited changes in UGP1 transcript levels under mannitol (−0.35) and salt (−0.15) treatments 24 h post stress initiation (Kilian et al., 2007); however, UGP1 transcripts are also known to be a poor indicator of protein-level or enzyme activity changes (Ciereszko et al., 2001).

UGP1 is an ubiquitous enzyme found in plants, animals, and bacteria that produces UDP-glucose (Meng et al., 2009), which is the direct substrate for trehalose synthesis (Schluepmann and Matthew, 2009). Thus, activation of UGP1 may be linked to increased trehalose biosynthesis in osmotic- and salt-stressed roots. In *E. coli*, trehalose stabilizes proteins and prevents protein aggregation during osmotic stress (Moruno Algara et al., 2019). This is also true for resurrection plants such as *Myrothamnus flabellifolius* where trehalose functions as a protectant to stabilize membranes and proteins, allowing them to survive during dehydration-rehydration cycles (Figueroa and Lunn, 2016; O’Hara et al., 2013). There is evidence that trehalose synthesis in Arabidopsis may serve a similar function during drought stress (Lin et al., 2019). Furthermore, knockdown of UGP1 in *S. cerevisiae* results in a reduction in trehalose as well as a reduction in oxidative stress resistance (Yi and Huh, 2015), suggesting that UGP1 involvement in mitigating osmotic stress is an evolutionary conserved mechanism.

In plants, stachyose and raffinose family oligosaccharides (RFO) are degraded by UGP1 (PWY-6527; https://biocyc.org/), releasing diphosphate. Thus an upregulation of UGP1 protein-levels and/or adjustments to its activity could serve as a homeostatic mechanism for adjusting the nutritional status of a plant under drought stress. Alternatively, if UGP1 lysine acetylation decreases functionality, it may increase RFO content in the cell. RFOs are suggested to have a dual role as osmoprotectants and compatible solutes (Elsayed et al., 2014). Other groups have found higher RFO content in desiccated plant tissues (Obendorf and Gorecki, 2012) and that RFOs stabilize membranes during dehydration (Pluskota et al., 2015). Overall, our data suggests that UGP1 functionality may be regulated by reversible lysine acetylation under both osmotic and salt stress conditions. However, whether UGP1 lysine acetylation increases or decreases UGP1 activity, and how that activity confers osmotic protection, is beyond the scope of this study.

### Lysine acetylation of GTP-Proteins may support root tip mitotic division

In Arabidopsis, GTP hydrolysis by Ran and Ran binding protein complexes are important in the regulation of auxin-induced mitotic progression in root tips (Kim and Roux, 2003; Xu et al., 2016). GTP hydrolysis by Ran facilitates protein import and release into the nucleus (Clarke and Zhang, 2008). Moreover, Ran facilitates mitotic spindle assembly by promoting the release of spindle assembly factors from importin complexes in the nucleus (Carazo-Salas et al., 1999). The role of Ran in mitotic spindle assembly has been proposed as an important role in root growth during drought stress (Akashi et al., 2016), and to intersect with ABA response and drought tolerance (Luo et al., 2013). In particular, both AtRAN1 and AtRAN3 are known to be induced by salt treatment, with *ran1/ran3* double mutants showing higher sensitivity to salt treatment, displaying significantly reduced root growth after being treated for 5 days with 100 mM NaCl (Xu et al., 2016).

Our data supports the involvement of RAN-GTPases in osmotic and salt stress responses. In particular, we find SIRANBP (AT1G07140) and RAN3 (AT5G55190) to be acetylated under salt (Log_2_FC = 1.92 SIRANBP; Log_2_FC = 3.68 AtRAN3) and mannitol (Log_2_FC = 1.81 SIRANBP; Log_2_FC = 3.65 RAN3) treatment (Figure 4–5). Phylogenetic analysis of RAN-GTPases Rab domains finds a 81 % (≤ 3 residue change) sequence homology between unicellular eukaryotes to vertebrates (Coppola et al., 2018), rendering it interesting that Ran GTPases in humans and mice are also regulated by reversible protein acetylation (De Boor et al., 2015). Human Ran possesses 5 lysine (K) acetylation sites: K37, K60, K71, K99, and K159 (Choudhary et al., 2009; PHOSIDA, http://www.phosida.com). Each site provides a different regulatory mechanism, for example, AcK159 increases HsRan binding towards Importin-β enhancing substrate import and release into the nucleus (De Boor et al. 2015). A global alignment between HsRan1 and AtRAN3 shows 81.6 % sequence similarity with HsK159 and AtK162 sharing an equivalent position (EMBL-EBI, Needle; Madeira et al., 2019). At the present moment, there are no studies reporting the effects of AcK162 on AtRAN3 in relation to osmotic or salt stress, however, considering the high sequence similarity between HsRan1 and AtRAN3 it is likely that AtRAN is regulated by lysine acetylation, and that this AcK162 may be responsible for increased mitotic division driven by RAN GTPase activity. Therefore, our observation of increased AtRAN-GTPase lysine acetylation may be a mechanism by which plants drive root elongation under drought stress conditions.

### ABF3 phosphorylation may aid in the intermediate and long term drought response

As a last resort, plants can launch the drought-escape (DE) response when tolerance or avoidance becomes an improbable solution to drought stress (Kooyers, 2015). DE is an adaptive mechanism that shifts the program from vegetative growth to reproductive growth triggering early flowering (Shavrukov et al., 2017). By shortening their life cycle they can set seed before the stress leads to their inevitable death. In rice, ABA promotes DE through positive regulation of flowering genes (Du et al., 2018). In Arabidopsis, the molecular mechanisms of DE are yet to be fully understood, nonetheless it is clear that under drought conditions Arabidopsis can favor early flowering (Kenney et al., 2014; Verslues and Juenger, 2011). Riboni et al. (2016) found ABA is capable of activating DE response by promoting transcriptional up-regulation of GIGANTEA (GI; AT1G22770) and florigen genes such as FLOWERING LOCUS T (FT; AT1G65480) and TWIN SISTER OF FT (TSF; AT4G20370) (Ribonu et al., 2016). A separate early flowering pathway involves ABA signaling through the expression and activity of ABSCISIC ACID RESPONSIVE ELEMENT-BINDING FACTOR 3/4 (ABF3; AT4G34000, ABF4; AT3G19290) transcription factors (Hwang et al., 2019). ABF3/4 interacts with the NF-Y complex to bind to the promoter of SUPPRESSOR OF CONSTANS 1 (SOC1; AT2G45660) and initiate early flowering. In our data, we find ABF3 phosphorylated as a result of salt and osmotic stress (NaCl_Log_2_FC = 3.89, MAN_Log_2_FC = 2.95). It is possible that ABF3 phosphorylation may promote the initiation of early flowering pathways needed for DE response. ABA induced multifaceted regulatory networks, such as florigen genes or direct activation of SOC1 transcription, may be necessary to ensure early flowering under drought-stress conditions.

Alternatively, up-regulation of ABF3 phosphorylation may be needed for drought avoidance (DA). ABFs also target the ABA-responsive regulatory *cis*-element (ABRE) and coordinate drought stress and ABA responses (Fernando et al., 2018; Liu et al., 2018). For instance, ABF3 directly regulates the expression of proteins, such as AtTPP1 (AT5G10100), which is required for the adjustment of stomatal aperture and primary root architecture (Lin et al., 2020, Kerr et al., 2018). ABA-dependent phosphorylation sites have been previously identified, with some sites conferring specificity to individual upstream kinases. For example, SNF1-RELATED KINASE 2 (SnRK2; AT3G50500) phosphorylates multiple conserved RXXS/T sites on ABF3 during ABA-dependent response to drought stress (Furihata et al. 2006; Uno et al. 2000; Yoshida et al., 2015). OPEN STOMATA 1 (OST1; AT4G33950) kinase phosphorylates ABF3 on multiple LXRXXpS/T preferred motifs including T451 (Sirichandra et al., 2010). As such, ABF3 controls part of the ABA-regulated transcriptome as an OST1 substrate. Calcium-dependent protein kinases 6 (CPK6; AT2G17290) also phosphorylates a subset of ABA-signaling related transcription factors, including ABF3 (Zhang et al., 2020). CPK6 phosphorylates ABF3 on preferred LXRXXpS/T motifs with the following sites identified: S32, S126, S134, and T169. However, we see ABF3 phosphorylated on S332, S338, and T342 (Supplemental Table 2), suggesting that these phosphorylation sites may be drought specific and the responsible protein kinases are unknown.

In the context of the 24 h stress presented in this study, it is more likely that an up-regulation of ABF3 phosphorylation is part of the plants DA response. Since ABF3 can be phosphorylated by multiple kinases, such as OST1 (Chen et al., 2013), CPK6 (Zhang et al., 2020), and SnKR2 (Fujii et al., 2009), it is probable that different phosphorylation events occur under DE and DA. The classification of DE and DA as discrete drought tolerance responses is unrealistic since a plant can employ a combination of DE and DA at different stages of its life cycle (Shavrukov et al., 2017). Thus it is probable that ABF3 phosphorylation is needed for both. Further research beyond the scope of this study is needed to identify the specific effect of these ABF3 phosphorylation events and the pathways that are initiated under DE and DA.

### Diverse PTM catalyzing enzymes exhibit abundance or phosphorylation status fluctuations in response to osmotic and salt stress

Our dataset uncovered a number of PTM modifying enzymes such as protein kinases (PKs; AT4G06744, AT5G48540, AT1G33590), phosphatases (AT3G45190) and ubiquitin ligases (AT2G23140) to change in overall abundance in response to either osmotic and/or salt stress. In particular, leucine rich-repeat (LRR) family protein AT4G06744, which decreases in abundance under both osmotic (Log_2_FC = −1.25) and salt stress (Log_2_FC = −3.73) and is exclusively root expressed (Klepikova Arabidopsis Atlas eFP browser - bar.utoronto.ca; Supplemental Table 1), but is otherwise uncharacterized. We also observe a decrease in the abundance of root development regulator PLANT U-BOX 4 (PUB4) an E3 ubiquitin ligase under salt treatment (Log_2_FC = −0.77), that when absent results in enhanced root growth (Kinoshita et al., 2015), suggesting that by 24 h, deregulation of root elongation has already been initiated at the protein level. Interestingly, the same authors suggest that PUB4 may intersect with auxin signaling to regulate root elongation, which would coincide with the extensive phosphorylation increases we observed on auxin related protein PILS2 (At1g71090; MAN_Log_2_FC = 4.21), TOM1-LIKE 4 (AT1G76970; MAN_Log_2_FC = 3.34, NaCl_Log_2_FC = 3.47) and AT1G79020 (MAN_Log_2_FC = 3.48, NaCl_Log_2_FC = 3.61) (Figure 5–6; Supplemental Table 2).

Our analysis of the osmotic and salt induced phosphoproteomes also revealed a number of protein kinases that possess osmotic and/or salt induced changes in their phosphorylation status, which aligns with our enrichment related biological processes (Supplemental Table 4). In particular, we observed 12 PKs and 1 ubiquitin ligase (AT1G22500) to change in their phosphorylation status under osmotic stress, while 17 PKs and 1 ubiquitin ligase (AT1G22500) changed in their phosphorylation status under salt stress, of which, only 5 are common between both stresses, indicating either stress has wide ranging impacts on a number of PKs and that there are distinct differences in the PKs impacted by each stress. With the majority of these PKs representing plasma membrane localized protein kinases that are uncharacterized it is likely that these represent key initiators of the phospho-signaling cascades that help determine ubiquitous and specialized responses to osmotic and/or salt stress. In particular, CANNOT RESPOND TO DMBQ 1 (CARD1 / HPCA1; AT5G49760; MAN_Log_2_FC = 3.54, NaCl_Log_2_FC = 2.45), which is phosphorylated under both osmotic and salt stress conditions and was recently resolved as a hydrogen peroxide sensor kinase (Wu et al., 2020), making it logical that the increase in phosphorylation is likely related to its activation under these stress regimes. This also renders the other uncharacterized PKs potentially exciting targets for further characterization.

Lastly, a number of previously characterized PKs were also resolved in our dataset as specific to either mannitol or salt treatment. Under mannitol treatment this includes: CALCIUM DEPENDENT PROTEIN KINASE 3 (CPK3; AT4G23650; Log_2_FC = 0.95), CAMODULIN-DOMAIN PROTEIN KINASE 9 (CPK9; AT3G20410; Log_2_FC = −0.75) and MAP3K EPSILON PROTEIN KINASE (MAP3KE1; AT3G13530; Log_2_FC = −1.14) (Figure 5; Supplemental Table 2) and CDK-ACTIVATING KINASE 4 (CAK4; AT1G66750; Log_2_FC = 1.25) under salt stress conditions (Figure 6; Supplemental Table 2). With increased H_2_O_2_ leading to increased intracellular Ca^2+^ (Wu et al., 2020), it is likely that CPK3 and CPK9, both exhibiting high levels of root expression (Klepikova Arabidopsis Atlas eFP browser - bar.utoronto.ca), represent important intermediaries in converting Ca^2+^ signaling responses to cellular outcomes. In particular, CPK3, which has been implicated in ABA promoted stomatal closure (Mori et al., 2006) and CPK9 which has a negative role in ABA-mediated stomatal closure (Chen et al., 2019). Our phosphoproteome data extend these opposing roles for CPK3 and 9 to root osmotic stress response through their opposing changes in phosphorylation status under osmotic stress (Figure 6; Supplemental Table 2). Lastly, we observe increased phosphorylation of CAK4 under salt stress. Given the central role of CAK4 in regulating cell division, and its expression in the root apical meristem, our data suggests that its phosphorylation likely promotes its cell division activity in order to drive root elongation under salt stress (Takatsuka et al 2009, Sofroni et al., 2020).

Overall, it is clear that protein phosphorylation and a subselection of protein kinases are particularly important in osmotic and salt stress response 24 h after initiation. Unlike PKs that may have exhibited transcriptional up-regulation, the subselection of PKs we observe here as changing in their abundance or phosphorylation status, represent excellent new candidates for future targeted characterization studies.

## CONCLUSION

Here, we describe proteomic and PTMomic changes occurring in Arabidopsis roots exposed to 300 mM mannitol and 150 mM NaCl for 24 h. By examining intermediate proteome responses, we have uncovered a number of intriguing candidate proteins and PTM events for future investigations, in addition to resolving new extended function for proteins with known roles in either osmotic or salt stress responses, and by extension, ABA signaling. that may be key to longer-term survival under osmotic and/or salt stress conditions. Furthermore, we show lysine acetylation to be a potentially interesting PTM in osmotic- or salt-stress response, with stress-induced changes having been particularly focused on key primary metabolic enzymes. In undertaking this quantitative analysis of the proteome, phosphoproteome and acetylome, we were able to capture a wide swath of the intermediate term protein responses in a single study. Moving forward, understanding the intermediate to long-term responses of either osmotic and/or salt stress at the protein-level through time-course experimentation, is especially important for the development of new climate resilient crops.

## MATERIALS & METHODS

### Plant Growth and Harvesting

*Arabidopsis thaliana* Col-0 seeds were imbibed 2 days at 4 °C in the dark prior to growth in Magenta boxes containing 0.5 x MS media containing 0.5 % (w/v) sucrose. Plants were grown at 22 °C under a 12 h light: 12 h dark photoperiod for 21 days prior to the application of 0.5 x MS media containing 300 mM mannitol (osmotic) or 150 mM NaCl (salt). Seedlings were grown in either experimental or control media for an additional 24 h, with stresses beginning and ending at zeitgeber time 3 (ZT 3). Roots were manually separated from shoots and immediately frozen in liquid N_2_. Roots were then ground in a mortar and pestle under liquid N_2_ and stored at −80 °C until further use.

### Sample Preparation and PTM Affinity Enrichments

Root tissue (800 mg) was extracted, processed and digested with trypsin (Promega) as previously described (Uhrig et al., 2019). Subsequent enrichment of phosphorylated or acetylated peptides was performed using titanium dioxide (NP-Ti02; Sachtopore; SNX 030S 005 #9205/1639) and anti-acetylated lysine IgG-coupled agarose (ICP0388; ImmuneChem, https://www.immunechem.com) as previously described (Uhrig et al., 2019), prior to analysis by mass spectrometry.

### Liquid Chromatography MS/MS

A total of 1 μg of re-suspended peptide was injected by an Easy-nLC 1000 system (Thermo Scientific) and separated on a Thermo Scientific™ EASY-Spray™ HPLC Column (75 μm x 500 mm; particle size 2.0 μm; E803A). The column was equilibrated with 100 % solvent A (0.1 % formic acid (FA) in water). Peptides were eluted using the following gradient of solvent B (0.1 % FA in ACN): 0-50 min; 0-25 % B, 50-60 min; 25-32 % B, 60-70 min; 32-98 % B at a flow rate of 0.3 μl/min. High accuracy mass spectra were acquired with an Orbitrap Fusion (Thermo Scientific) that was operated in data dependent acquisition mode. All precursor signals were recorded in the Orbitrap using quadrupole transmission in the mass range of 300-1500 m/z. Spectra were recorded with a resolution of 120 000 at 190 m/z, a target value of 4E5 and the maximum cycle time was set to 3 seconds. Data dependent MS/MS were recorded in the linear ion trap using quadrupole isolation with a window of 2 Da and HCD fragmentation with 30 % fragmentation energy. The ion trap was operated in rapid scan mode with a target value of 3E4 and a maximum injection time of 100 ms. Precursor signals were selected for fragmentation with a charge state from +2 to +7 and a signal intensity of at least 1E4. A dynamic exclusion list was used for 30 seconds and maximum parallelizing ion injections was activated.

### Mass Spectrometry Data Analysis

Raw data were processed using MaxQuant software version 1.6.14.0 (http://www.maxquant.org/; Cox and Mann, 2008) and searched against The Arabidopsis Information Resource (TAIR10) protein database concatenated with a decoy database supplemented with contaminants using the Andromeda search engine. Fixed modifications included carbamidomethylation of cysteine residues, while methionine oxidation (all searches), lysine acetylation (acetylated peptide enrichments) and phosphorylated serine / threonine and tyrosine (phosphopeptide enrichments) were set as variable modifications, respectively. One missed cleavage for phosphoproteomic and proteomic measurements and two missed cleavages for lysine-acetylated peptides were permitted. A protein and PSM false discovery rates (FDR) threshold of 1 % was used, while match between runs and re-quantify options were enabled. Further downstream analysis was performed using Perseus version 1.6.14.0 (Tyanova et al., 2016). This involved the removal of reverse hits and contaminants, followed by log_2_-transformation, assembly into treatment groups and filtering based on the presence of measured data in at least 2 replicates of at least one group. Data were then median normalized, followed by imputation of missing values based on the normal distribution function set to default parameters. All PTM analyses utilized a PTM site localization score threshold ≥ 0.75. Significant changes in either protein or PTM peptide levels between each treatment condition and the control samples were assessed by Student’s T-test (*pval* ≤ 0.05). Mass spectrometry data acquired for this study can be found at the EMBL proteomic repository Proteomics IDEntifications (PRIDE; https://www.ebi.ac.uk/pride/archive/; PXD019139).

### Bioinformatics

To identify the biological functions of proteins changing in abundance, phosphorylation or lysine acetylation, a gene ontology (GO) analysis of biological processes was performed using the ontologizer (http://ontologizer.de; (Bauer et al., 2008). A parent–child intersection analysis approach was used (Benjamini–Hochberg *FDR* correction *pval* ≤ 0.05). Clustering of Arabidopsis microarray time-course data was performed using a trajectory-based longitudinal clustering algorithm implemented in the kml-shape package in R (Genolini et al., 2016). For inputs, microarray counts were obtained from (Kilian et al., 2007) and fold changes were calculated for each time point comparing treated samples versus the controls. The resulting dataset was filtered based on the set of proteins that were found to significantly change in either abundance, phosphorylation or lysine acetylation status at 24 h. The resulting transcript fold-changes over the time-course were used to create four clusters based on the transcript trajectories over the 1, 3, 6, 12 and 24 h time points previously obtained (Kilian et al., 2007). Clustered trajectories and corresponding protein data were plotted using the *ggplot2* and *cowplot* packages in R version 3.3.1 (R Core Team 2016). Further characterization of connections between quantified and significantly changing phospho- and acetyl-proteins was performed through association network analyses using the Cytoscape StringDB plugin StringApp (http://apps.cytoscape.org/apps/stringapp; Szklarczyk et al., 2017) with an overallStringDB association score threshold of 0.8. Cytoscape 3.8.2 (http://www.cytoscape.org/) visualized networks were then supplemented with subcellular localization data obtained from SUBAcon (http://suba.live/; (Hooper et al., 2014)

## ACKNOWLEDGEMENT

The authors would like to thank the Functional Genomics Center Zurich for helpful discussions and the National Science and Engineering Research Council of Canada (NSERC) for funding this research.

## CONFLICTS OF INTEREST

The authors declare they have no conflicts of interest

## SUPPLEMENTAL INFORMATION

Supplemental Table 1: Results of the proteome analysis

Supplemental Table 2: Results of the phosphoproteome analysis

Supplemental Table 3: Results of acetylome analysis

Supplemental Table 4: Gene Ontology and protein domain enrichment for proteins with significantly changing abundance upon osmotic and salt stress.

Supplemental Table 5: Transcript data clustering analysis

## Notes

### Competing Interest Statement

The authors have declared no competing interest.

### Summary of Updates

In our original manuscript, we found the protein ARO2 to be down-regulated in abundance upon salt stress. We then aimed to corroborate this finding by assessing the phenotype of aro2-1; a TDNA insertional mutant under salt stress versus wild-type plants. However, upon re-performing the proteomic analysis, which required a newer version of MaxQuant, ARO2 was found to no longer pass significance thresholds. Overall, the dataset was found to be vastly similar to the original manuscript in terms of proteins exhibiting changes in the acetylation, phosphorylation and abundance. Given this, we have completely removed reference to ARO2 from the updated manuscript; however, despite the removal of this data, we believe the original phenotypic data we observed and originally reported for aro2-1 in relation to salt stress (which was from n = 100 seedlings) is correct.

